# Transcriptome-wide Cas13 guide RNA design for model organisms and viral RNA pathogens

**DOI:** 10.1101/2020.08.20.259762

**Authors:** Xinyi Guo, Hans-Hermann Wessels, Alejandro Méndez-Mancilla, Daniel Haro, Neville E. Sanjana

## Abstract

CRISPR-Cas13 mediates robust transcript knockdown in human cells through direct RNA targeting. Compared to DNA-targeting CRISPR enzymes like Cas9, RNA targeting by Cas13 is transcript- and strand-specific: It can distinguish and specifically knock-down processed transcripts, alternatively spliced isoforms and overlapping genes, all of which frequently serve different functions. Previously, we identified optimal design rules for *Rfx*Cas13d guide RNAs (gRNAs), and developed a computational model to predict gRNA efficacy for all human protein-coding genes. However, there is a growing interest to target other types of transcripts, such as noncoding RNAs (ncRNAs) or viral RNAs, and to target transcripts in other commonly-used organisms. Here, we predicted relative Cas13-driven knock-down for gRNAs targeting messenger RNAs and ncRNAs in six model organisms (human, mouse, zebrafish, fly, nematode and flowering plants) and four abundant RNA virus families (SARS-CoV-2, HIV-1, H1N1 influenza and MERS). To allow for more flexible gRNA efficacy prediction, we also developed a web-based application to predict optimal gRNAs for any RNA target entered by the user. Given the lack of Cas13 guide design tools, we anticipate this resource will facilitate CRISPR-Cas13 RNA targeting in common model organisms, emerging viral threats to human health, and novel RNA targets.

---

CRISPR-Cas13 mediates robust transcript knockdown in human cells through direct RNA targeting^1-4^. Compared to DNA-targeting CRISPR enzymes like Cas9, RNA targeting by Cas13 is transcript- and strand-specific: It can distinguish and specifically knock-down processed transcripts, alternatively spliced isoforms and overlapping genes, all of which frequently serve different functions. Previously, we have described a set of optimal design rules for *Rfx*Cas13d guide RNAs (gRNAs), and developed a computational model to predict gRNA efficacy for all human protein-coding genes^5^. However, there is a growing interest to target other types of transcripts, such as noncoding RNAs (ncRNAs)^6,7^ or viral RNAs^8,9^, and to target transcripts in other commonly-used organisms^10-13^. Here, we predicted relative Cas13-driven knock-down for gRNAs targeting messenger RNAs and ncRNAs in six model organisms (human, mouse, zebrafish, fly, nematode and flowering plants) and four abundant RNA virus families (SARS-CoV-2, HIV-1, H1N1 influenza and MERS). To allow for more flexible gRNA efficacy prediction, we also developed a web-based application to predict optimal Cas13d guide RNAs for any RNA target entered by the user.

To select optimal gRNAs for transcripts produced from the reference genomes of human, mouse, zebrafish, fly, nematode and flowering plants, we created a user-friendly Cas13 online platform (https://cas13design.nygenome.org/) (**Fig. 1a**). We previously found that optimal Cas13 gRNAs depend on specific sequence and structural features, including position-based nucleotide preferences in the gRNA and the predicted folding energy (secondary structure) of the combined direct repeat plus gRNA^5^. Using this algorithm, we pre-computed gRNA efficacies, where possible, for all mRNAs and ncRNAs with varying transcript length for the 6 model organisms (**Fig. 1b, Supplementary Fig. 1**).

**Figure 1.**
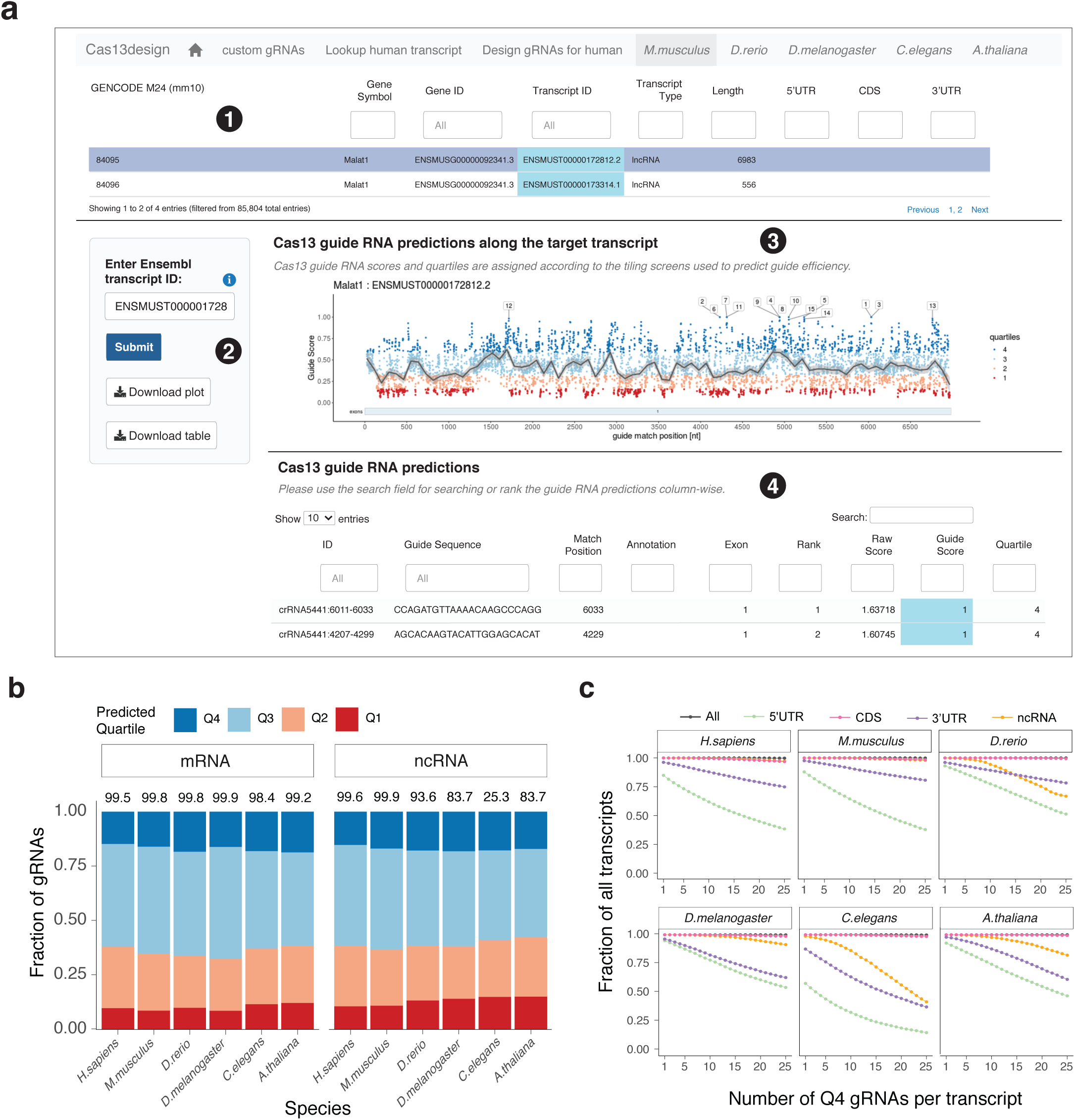
A graphical interface for optimized CRISPR-Cas13d gRNA design for messenger RNAs (mRNAs) and noncoding RNAs (ncRNAs) from six common model organisms. (**a**) Example output of Cas13 Design webtool. Circle (1) allows searches by gene symbol or transcript ID for gRNA design (2), with options to download generated plots and data tables. Circle (3) displays a distribution of gRNAs along the target transcript color-coded by the predicted targeting efficacy scores separated into four quartiles. Q4 gRNAs correspond to those with the highest predicted efficacy and Q1 correspond to those with the lowest predicted efficacy. Circle (4) displays sgRNA options with on-target score predictions. (**b**) The predicted guide efficacy quartiles for mRNAs and ncRNAs across six model organisms. Numbers above bars indicate the percent of transcripts scored that meet the minimal length requirement for target RNAs (80 nt). (**c**) The fraction of processed transcripts that contain at least 1 (up to 25) Q4 gRNAs (predicted high-scoring gRNAs).

For the scored gRNAs for each organism, we found that approximately 20% are ranked in the top quartile (Q4 guides) for both mRNAs and ncRNAs (**Fig. 1b**). Remarkably, even though the nucleotide composition can very between RNAs from different species^14-16^, we find a similar proportion of optimal *Rfx*Cas13d gRNAs across all six species.

Next, we examined how many predicted high efficacy gRNAs are present, on average, in different transcripts. To do this, we determined what fraction of the transcripts in each organism include *n* top-scoring (Q4) gRNAs for values of *n* between 1 and 25. We found that coding sequences contained a higher number of top-scoring gRNA per transcript across all organisms, whereas targeting the noncoding transcriptome is more challenging and varies across different organisms (**Fig. 1c**). On average, we were able to find at least 25 Q4 gRNAs for >99% of coding exons in mRNAs but only 80% of ncRNAs. Beyond targeting transcripts from the reference genomes of these model organisms, there are also many other applications of Cas13, such as targeting transcripts from non-model organisms, cleavage of synthetic RNAs, and targeting of transcripts carrying genetic variants not found in the reference genome. Therefore, in addition to these pre-scored gRNAs, we have also developed a graphical interface that allows the user to input a custom RNA sequence for scoring and selection of optimal gRNAs.

Recently, several groups have proposed using CRISPR-Cas13 nucleases to directly target viral RNAs^8,17^, which has become an area of rapid technology development due to the recent COVID-19 pandemic^18^. However, these approaches do not use optimized Cas13 guide RNAs. Previously, we showed that optimal guide RNAs targeting an EGFP transgene can result in a ∼10-fold increase in knock-down efficacy compared to other gRNAs^5^. Therefore, to speed development of effective CRISPR-based antiviral therapeutics, we applied our design algorithm to target SARS-CoV-2 and other serious viral threats using Cas13d.

To ensure coverage of diverse patient isolates, we collected 7,630 sequenced SARS-CoV-2 genomes submitted to the Global Initiative on Sharing All Influenza Data (GISAID) database from 58 countries/regions^19^ (**Fig. 2a**). Using the first sequenced SARS-CoV-2 isolate from New York City (USA/NY1-PV08001/2020) as a reference^20^, we evaluated how many individual SARS-CoV-2 genomes each reference gRNA can target (**Fig. 2b**). Guide RNAs targeting protein-coding regions are mostly well-conserved across all genomes, with lower conservation in more variable regions such as Non-Structural-Protein 14 (NSP14) and Spike (S) protein. We found that gRNAs targeting in the 5’ and 3’ untranslated regions tended to be poorly conserved, as might be expected given the lack of coding function of these regions (**Supplementary Fig. 2**). Upon examination of each of the 26 SARS-CoV-2 genes, we found that all gene transcripts could be targeted with Q4 gRNAs.

**Figure 2.**
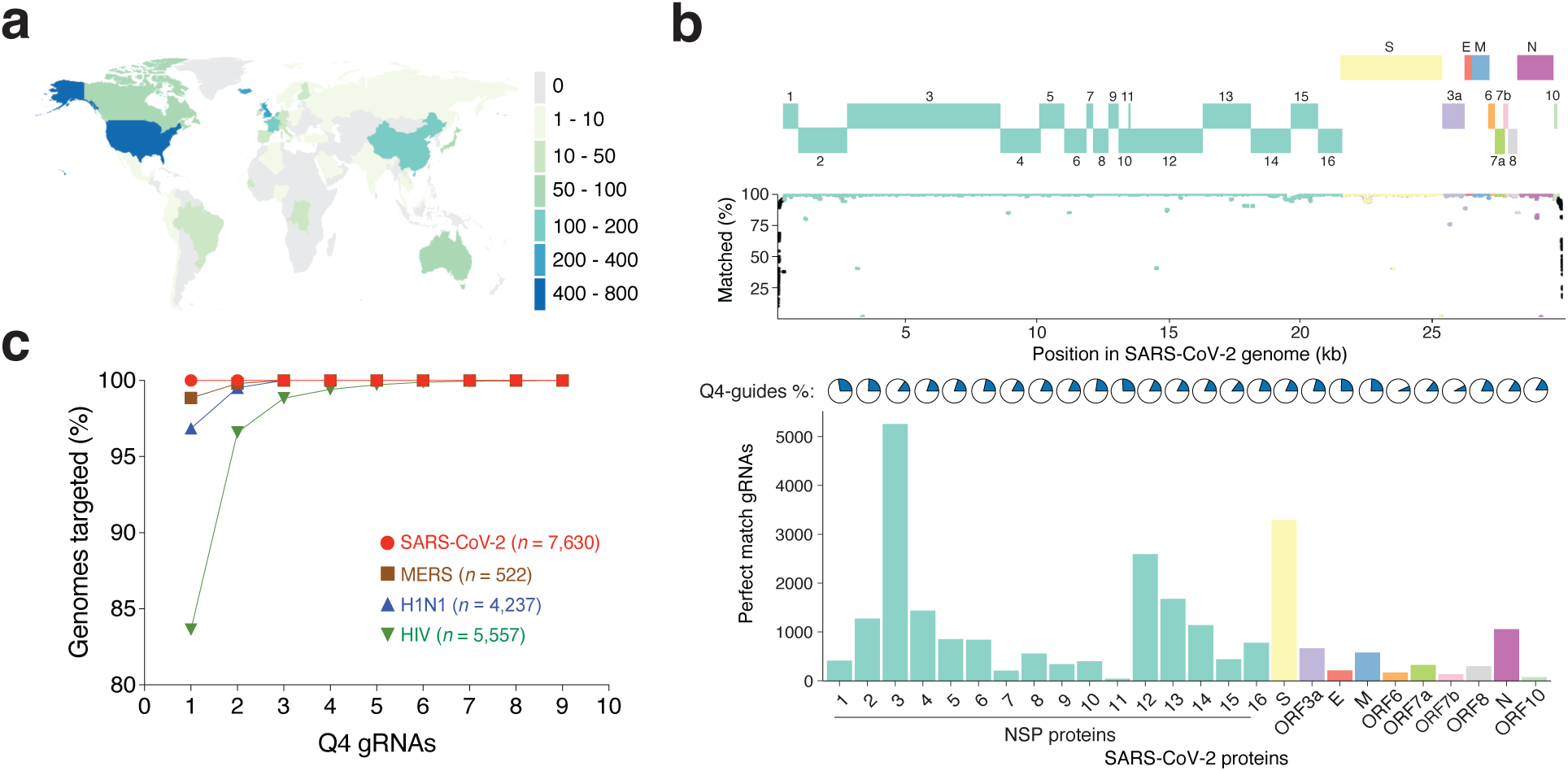
Optimal CRISPR-Cas13d gRNAs to target common human pathogenic RNA viruses. **(a)** World map of analyzed SARS-CoV-2 isolates (data from GISAID, April 17^th^, 2020). **(b)** Guide RNA design for each SARS-CoV-2 gene. *Top panel:* SARS-CoV-2 gene annotations. *Middle panel:* Percent of SARS-CoV-2 genomes targeted by each NY1 reference gRNA. *Bottom panel*: Fraction of gRNAs in Q4 per gene (pie) and total number of Q4 gRNAs per gene that targets at least 99% of the total genomes (bar). **(c)** Predicted minimum number of Q4 gRNAs to target all SARS-CoV-2, MERS-CoV, H1N1, and HIV-1 genomes analyzed (*n* = 7630, 522, 4237 and 5557 viral genomes, respectively).

Similarly, we designed and scored all gRNAs for the coronavirus MERS and two other RNA viruses, HIV-1 which drives Acquired Immunodeficiency Syndrome (AIDS) and H1N1 pandemic influenza. Unlike SARS-CoV-2, where a single high-efficacy (Q4) gRNA can target all genomes analyzed, we found that at least two gRNAs are needed to target nearly all available genomes. For the highly mutagenic virus HIV-1^21^, we found that nine gRNAs are needed to target all available genomes (**Fig. 2c**). Given the tremendous interest in viral RNA targeting using Cas13 enzymes, this dataset of optimized gRNAs provides a platform for the development of CRISPR therapeutics for broad targeting of viral populations from diverse patient isolates. All designed gRNAs for model organism and viral transcripts can be interactively browsed or downloaded in bulk on the design tool website.

RNA-targeting CRISPR-Cas13 has great potential for transcriptome perturbation and antiviral therapeutics. In this study, we have designed and scored Cas13d gRNAs for both mRNAs and ncRNAs in six common model organisms and identified optimized gRNAs to target virtually all sequenced viral RNAs for SARS-CoV-2, HIV-1, H1N1 influenza and MERS. We further expanded our web-based platform to make the Cas13 gRNA design readily accessible for model organisms and created a new application to enable gRNA prediction for user-provided target RNA sequences. Given the current lack of Cas13 guide design tools, we anticipate this resource will greatly facilitate CRISPR-Cas13 RNA targeting in model organisms, emerging viral threats to human health and novel RNA targets.

## Acknowledgements

We thank the entire Sanjana laboratory for support and advice. We thank M. Zaran and S. Brock for assistance with the web-tool server. N.E.S. is supported by New York University and New York Genome Center startup funds, National Institutes of Health (NIH)/National Human Genome Research Institute (grant nos. DP2HG010099), NIH/National Cancer Institute (grant no. R01CA218668), Defense Advanced Research Projects Agency (grant no. D18AP00053), the Sidney Kimmel Foundation, the Melanoma Research Alliance, and the Brain and Behavior Foundation.

## Author contributions

N.E.S. and H.H.W. conceived the project. N.E.S., H.H.W., X.G. and A.M.-M. designed the study. A.M.-M. and D.H. designed Cas13d gRNAs for all model organisms. H.H.W built the web tool and performed analyses for model organisms. X.G. designed Cas13d gRNAs and performed analyses for viruses. X.G., H.H.W and D.H. produced the figures. N.E.S. supervised the work. All authors contributed to drafting and reviewing the manuscript, provided feedback and approved the manuscript in its final form.

## Methods

### gRNA design for model organisms

Reference transcriptomes and corresponding annotations were obtained for each model organism: *H. sapiens* (GENCODE v19, GRCh37), *M. musculus* (GENCODE M24, mm10), *D. rerio* (Ensembl v99, GRCz11), *D. melanogaster* (Ensembl v99, BDGP6), *C. elegans* (Ensembl v99, WBcel235) and *A. thaliana* (Ensembl Plants v46, TAIR10). For each organism, we performed the on-target efficiency predictions for both mRNAs and ncRNAs using command-line *Rfx*Cas13d designer version 0.2 as previously described^5^. We scored gRNAs for all RNA targets with a length of at least 80 nucleotides.

### RNA virus genome collection

All full-length RNA virus genomes were downloaded on April 17^th^, 2020. We downloaded 7,630 complete SARS-CoV-2 viral genomes classified as high coverage and 4,237 Influenza A H1N1 viral genomes with a complete set of eight genomic segments. SARS-CoV-2 and H1N1 genomes were obtained from GISAID (https://www.gisaid.org/). We also analyzed 522 MERS-CoV and 5,557 full length HIV-1 viral genomes, which were downloaded from NCBI Virus (https://www.ncbi.nlm.nih.gov/labs/virus/).

### gRNA design to target SARS-CoV-2

We split multi-FASTA files into single-entry FASTA files using the UCSC tool faSplit^22^. All possible 23-mer gRNAs targeting individual genomes were scored with the *Rfx*Cas13 on-target model described previously^5^. All scored guides were classified into four quartiles. Quartile 4 guides (or Q4) are designated to be the predicted best-performing guides. We used USA/NY1-PV08001/2020 (refer to as NY1 isolate) for the SARS-CoV-2 reference gRNA design. Compared to the original (Wuhan) isolate, NY1 contains 3 nucleotide substitutions (G3243A, C25214T, G29027T) resulting in two amino acid mutations (N: A252S, ORF1a: G993S). The SARS-CoV-2 transcript annotation was obtained from NCBI (GenBank: NC_045512.2).

### Prediction of minimal numbers of gRNAs to target RNA viruses

For each RNA virus, we identified a minimal set of high-scoring Q4 gRNAs that could target all genomes collected. We used a greedy algorithm as described previously^8^: For each iteration, the gRNAs with the highest number of targeting genomes are added to the set. During each iteration, if multiple gRNAs target the same highest number of genomes, we will pick one for the minimal set and start the next iteration.

### Code availability

All designed Cas13 guide RNAs (for model organisms and RNA viruses) and the interactive design tool are available here: https://cas13design.nygenome.org/. For additional reproducibility, we provide shell scripts and R code to reproduce the figures here: https://gitlab.com/sanjanalab/cas13_webtool. The Cas13 guide design algorithm is available here: https://gitlab.com/sanjanalab/cas13.

