## Supplementary Figures for "Transcriptome-wide Cas13 guide RNA design for model organisms and viral RNA pathogens"

**Supplementary Figure 1. Transcript length for mRNAs and ncRNAs across species.**

**Supplementary Figure 2. Q4 gRNAs targeting coding SARS-CoV-2 regions verses noncoding SARS-CoV-2 regions.**

**Supplementary Figure 1**

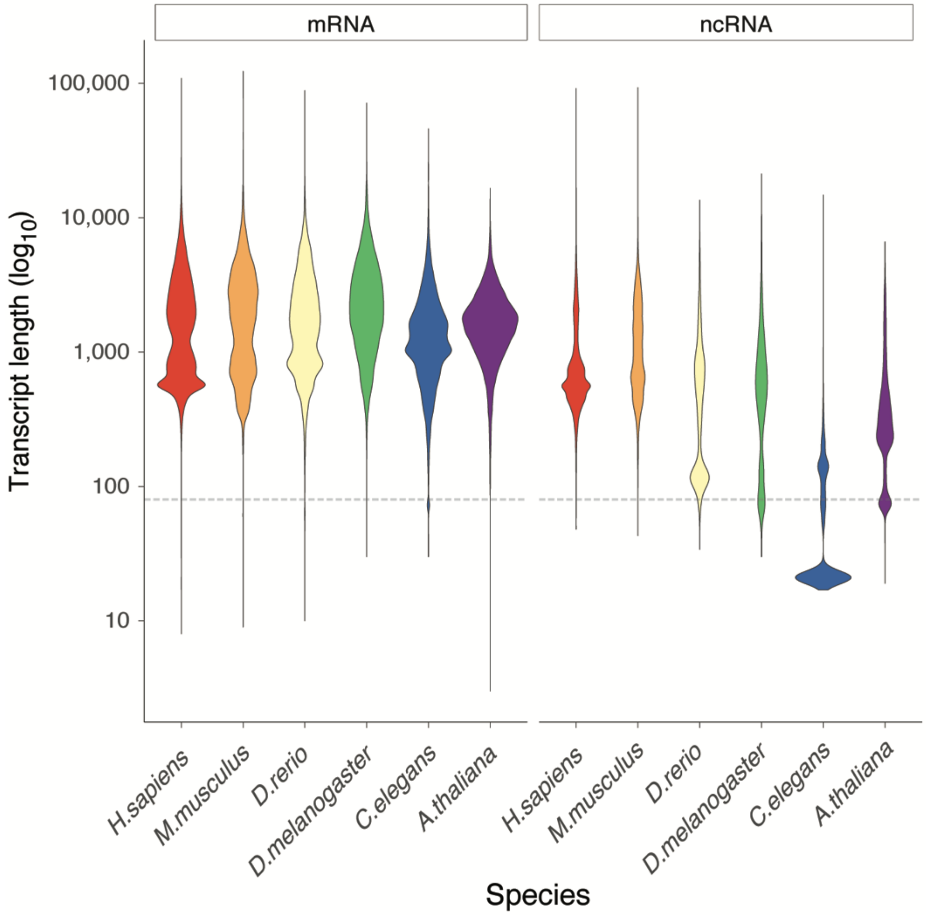


**Supplementary Figure 1. Transcript length for mRNAs and ncRNAs across species.** Dotted line indicates the minimal input length requirements (> 80 bp) for Cas13d design software. Transcript lengths were derived from corresponding gene annotation reference sequences.

**Supplementary Figure 2**

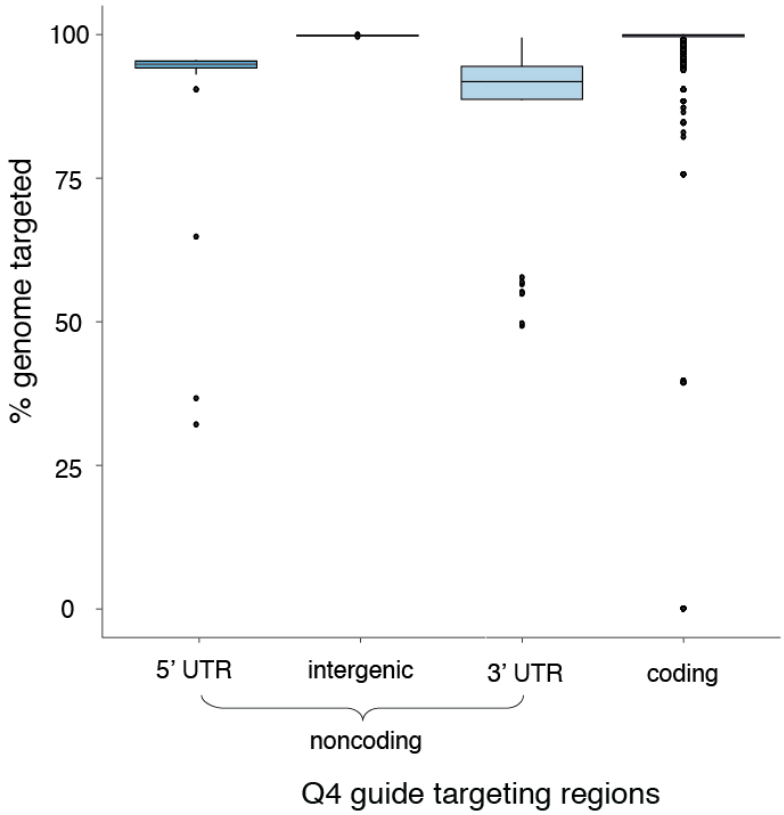


**Supplementary Figure 2. Q4 gRNAs targeting coding SARS-CoV-2 regions verses noncoding SARS-CoV-2 regions.** Classification of coding and noncoding regions is from the NCBI annotation of the SARS-CoV-2 reference strain (see *Methods*).
